# Layered Feedback Control Improves Robust Functionality across Heterogeneous Cell Populations

**DOI:** 10.1101/2020.03.24.006528

**Authors:** Xinying Ren, Richard M. Murray

## Abstract

Realizing homeostatic control of metabolites or proteins is one of the key goals of synthetic circuits. However, if control is only implemented internally in individual cells, cell-cell heterogeneity may break the homeostasis on population level since cells do not contribute equally to the production or regulation. New control structures are needed to achieve robust functionality in heterogeneous cell populations. Quorum sensing (QS) serves as a collective mechanism by releasing and sensing small and diffusible signaling molecules for group decision-making. We propose a layered feedback control structure that includes a global controller using quorum sensing and a local controller via internal signal-receptor systems. We demonstrate with modeling and simulation that the global controller drives contributing cells to compensate for disturbances while the local controller governs the fail-mode performance in non-contributing cells. The layered controller can tolerate a higher portion of non-contributing cells or longer generations of mutant cells while maintaining metabolites or proteins level within a small error range, compared with only internal feedback control. We further discuss the potential of such layered structures in robust control of cell population size, population fraction and other population-dependent functions.

## I. INTRODUCTION

In synthetic biology, one important challenge is to maintain homeostasis from single-cell level to large-scale multicellular systems using proper control. Negative feedback is an essential strategy for such processes that requires sensing disturbances and adapting [1], [2]. Much research has focused on implementing feedback controllers that robustly regulate metabolites or protein concentrations on single-cell level. Two general implementations of negative feedback are inhibiting protein production and enhancing protein degradation [3]–[5], and they are widely used in regulating metabolic biosynthesis [6], biofuel production [7] and dose-response [8]. Integral negative feedback is an appealing controller since it effectively drives the regulated protein level to a constant set-point without error [9]. Using a strong sequestration pair of a *σ* -factor and a anti-*σ* -factor, an antithetic integral feedback controller has been recently implemented for robust perfect adaptation [10].

However, internal feedback control on single-cell level does not always lead to population level homeostasis. Cell-cell heterogeneity is commonly observed in bacteria, yeast and mammalian cell communities [11]–[13]. Diverse phenotypes and behaviors help organisms to adapt to fluctuating environments and to better survive as a bet-hedging strategy [14], [15]. For example, the persistence mechanism in *E. coli* allows some cells exhibiting the persistent state and prolongs the population’s survival when exposed to antibiotics [16]. Mutation is another source of population heterogeneity and often cheaters gain more benefits without paying costs [17], [18]. In biofilm formation, individual cells follow different developmental pathways that also leads to heterogeneous populations [19]. Non-contributing cells, i.e., cells that are switched to a different state under stress or cheater cells, no longer perform identically as the contributing cells, since the production or regulation may be far off expected. Under these conditions, population level homeostasis of metabolites or proteins is significantly perturbed when these non-contributing cells take a larger fraction of the whole population after generations, and the internal controller cannot respond to such disturbances properly.

Population level homeostasis requires more control structures on top of the internal feedback. Quorum sensing systems are commonly observed in bacteria for sensing the collective behaviors across the whole population and directing responses in individual cells [20], and have been used in synthetic circuits to engineer microbial consortia [21]–[23]. A typical quorum sensing system utilizes diffusible AHL molecules mediated by the LuxI-LuxR families. LuxI proteins governs AHL sythesis and LuxR proteins are AHL-triggered receptors that regulate downstream transcriptions [24]. Since AHL molecules diffuse across membranes and well mix in the environment, the global AHL concentration is often regarded as a measurement of populational bahaviors. Therefore, we can build a global feedback controller where the target protein regulates the AHL synthase LuxI and AHL-triggered receptors regulate the transcription of the target protein. When a heterogeneous population appears, the global feedback controller in contributing cells can detect variations on population level behaviors and respond properly via AHLs. Meanwhile, we can rewire relationships between the target protein and LuxR proteins to build a local feedback so that non-contributing cells apply different control actions than contributing cells to improve fail-mode performance and thus maintain populational homeostasis.

We demonstrate such a layered controller with global and local feedback via quorum sensing signals and receptors improves robustness in following sections. In Section II, we introduce a simple protein regulation circuit with internal feedback control using repressors. In Section III, we show how to build a global controller and a layered controller based on the same repressing control law in the internal controller. In Section IV, we show mathematical analysis and simulations of the internal, global and layered controller performance and compare the steady state error of the protein level in heterogeneous populations. In Section V, we further discuss potential applications of the layered controller in more population control problems.

## II. Internal Feedback Control on Protein Level

We consider a protein regulation circuit in *E. coli* shown in Fig. 1(a). We introduce a simple feedback where the target protein’s transcription is repressed by a repressor and the production of the repressor is activated by the target protein. When the protein concentration is perturbed to a higher/lower level, more/fewer repressors get produced and weaken/strengthen the protein transcription, thus the closed-loop keeps a constant protein concentration in individual cells. We develop an ODE model to characterize the closed-loop dynamics:

**Fig. 1.**
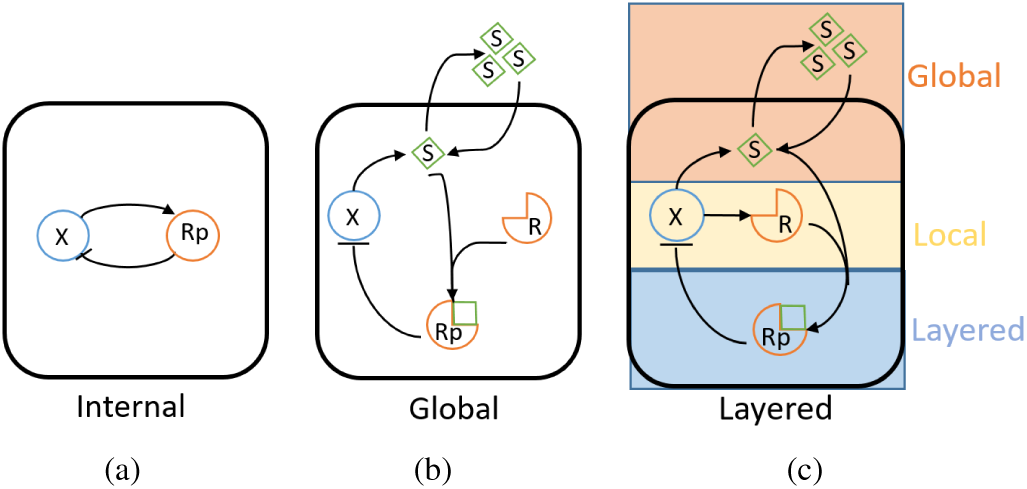
Sketches of protein regulation circuits using different controllers. Panel (a) shows the internal feedback on the target protein *X* via a repressor *Rp*. In (a), *X* activates *Rp* and *Rp* represses transcription of *X* to form a negative feedback loop. Panel (b) implements the global feedback. In (b), *X* activates the synthesis of diffusible AHL molecules *S. S* can bind with a constitutive receptor *R* to trigger the repression on *X* via *Rp*. Panel (c) demonstrates a layered feedback with a global controller as in panel (b) and a local controller where *X* also activates *R*.

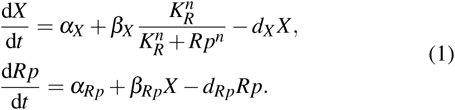

In equation (1), *X* is the target protein with a constitutive production and a Hill-type repression by the repressor *Rp*. We assume *X* activates the transcription of *Rp* in the linear regime and *Rp* serves as a proportional control to *X*. Both *X* and *Rp* dilute with cell division. Assuming that the activation in *Rp* transcription is inducible, the feedback strength can be tuned to set the steady state level of *X* by altering the rate *β*_*Rp*_.

To show that the protein level in individual cells can be regulated and maintains a stable steady state with the internal feedback controller, we simulate the dynamics when intracellular fluctuations affect the protein concentration. In Fig. 2(a), we alter the induction level of the repressor from low to high(colored from light green to dark green) and the protein levels converge to different values. At time *t* = 120 min, the protein level is perturbed and the internal feedback recovers the steady states. Assuming that in homogeneous populations, individual cells perform identically, then the protein dynamics shown in Fig. 2(a) should also represent the population level protein dynamics. The repressed production kinetics of the protein is usually assumed to follow the Hill function to provide reasonable tunabililty and sensitivity [25]. In Fig. 2(b), we show the open-loop relationship between the repressor and the protein level(colored blue) and the internal feedback with different strength(colored green). The intersection points of the open-loop and the feedback control curves are equilibria that the protein is expected to converge to at steady state. We consider the regimes within the gray dashed lines as an ideal working regime of the circuit.

**Fig. 2.**
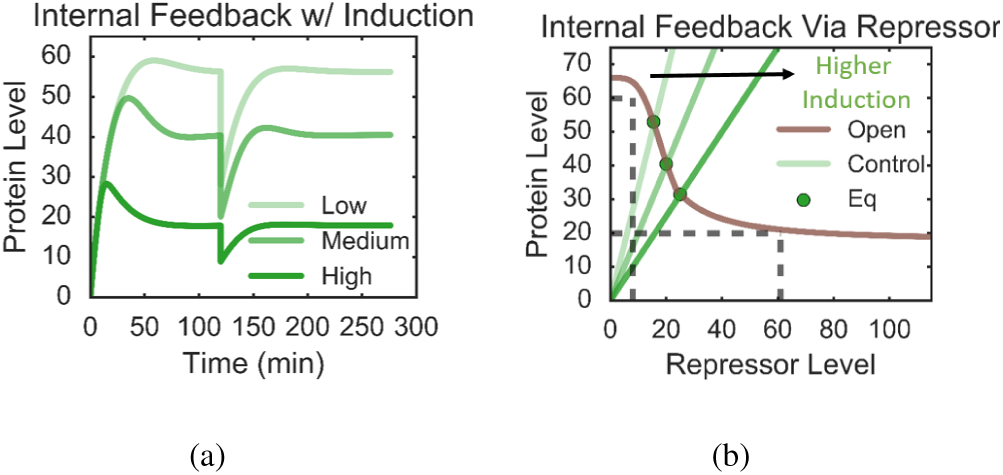
Simulations and tuning response curves of the internal feedback circuit. Panel (a) shows that the protein level can be set to different constant values with inductions on the repressor. When the protein is perturbed to a lower concentration, the internal feedback recovers its previous level. Panel (b) is the tuning curve of the target protein by the repressor. The control curve determines the equilibrium and shows that *X* is tunable within a range of *Rp*.

## III. Layered Feedback Control on Protein Level

The layered controller includes a quorum sensing system to trigger the repressor [26]. Both the signaling molecules and the receptors are regulated by the target protein and can bind to form a complex functioning as a repressor. Instead of the direct feedback from the intracellular protein to the repressor, the layered feedback can sense population behaviors via signaling molecules AHLs and individual cell behaviors via receptors LuxR at the same time before applying actuation on protein production through the triggered repressor. The key principle to realize such layered controllers is to have separate global feedback via the signaling molecules and the local feedback via receptors. Therefore the control action through the signal-receptor complex carries information of both the whole population and individual cells.

Similar strategies using quorum sensing systems for population control have been proposed and implemented in previous studies [27]–[30]. Most of these circuits are designed to express a constitutive receptor and only regulate AHL synthesis as a global feedback. By choosing the working regime, the triggered activator or repressor approximately depends on AHL level linearly. Consider that a single AHL molecule *S* and a single receptor *R* bind to form a complex *Rp* with binding rate *k*^+^ and unbinding rate *k*^−^, the complex level at steady state is determined by the Michaelis-Menten equation with the dissociation constant 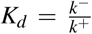 as the following:

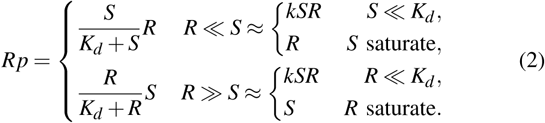

When *R*≪*S, Rp* rises approximately linearly with *S* if *S*≪*K* and reaches saturation when *S* is high. Similarly, we can approximate *Rp* in a linear regime and a saturation regime when *R*≫*S*.

Now we show how to achieve a global controller with a constitutive receptor and a layered controller where both AHL and receptor expressions are regulated within multiple potential regimes.

### A. Global Feedback Control with A Constitutive Receptor

To build a global feedback where cells sense AHLs to trigger protein repression, we include a constitutive receptor as shown in Fig. 1(b) and ensure the triggered repressor depends linearly on AHL concentration. According to equation (2), one possible design is to express more receptors than AHLs. Another design is to express a low level of receptors and keep the range of AHLs lower than the dissociation constant *K*_*d*_, which suggests choosing a weak binding signal-receptor binding pair for a large linear working regime.

We consider the second scenario where *R* ≪ *S*. Given the following assumptions, we obtain a simplified model of the target protein *X*, the intracellular AHL *S*, the triggered repressor *Rp*, and the global AHL 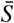: 1) total cell population size is fixed as *N*; 2) AHL molecules diffuse in and out membranes at the same rate *D*_*f*_ and diffuse freely and get mixed quickly in the environment [31]; 3) the signal-receptor binding reaction is fast compared to transcription and translation and can be characterized as a Michaelis-Menten equation. This yields a model:

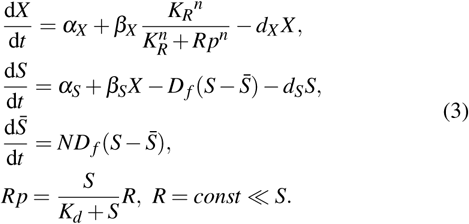

In homogeneous populations, the global feedback behaves the same way as the internal feedback described by equation (1) before AHLs reach saturation. By altering the induction on AHL synthesis, the global feedback also presents a similar tunability, as shown in Fig. 3(a). To better illustrate how the approximations of equation (2) determine different working regimes, we plot both AHL and triggered repressor levels and show they diverge when switching from linear to saturating regimes in Fig. 3(b).

**Fig. 3.**
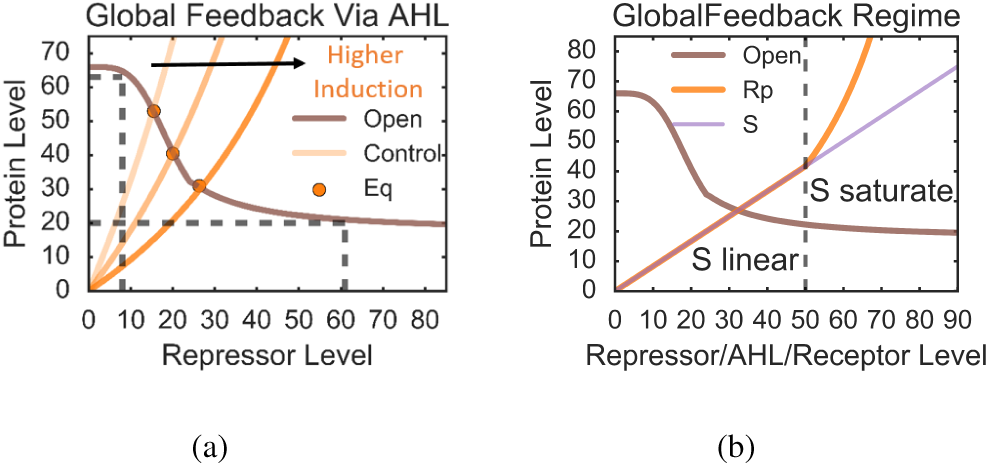
Tuning response curves of the global feedback circuit. Panel (a) shows that the target protein level can be tuned by inducing the AHL synthesis at different strengths. The control actuation shows saturation in the triggered repressor when AHL concentration is high. Panel (b) demonstrates a more detailed switch in working regimes from linear to saturation when AHL level rises above the threshold.

### B. Layered Feedback Control

The layered controller involves a global feedback via AHLs and a local feedback via receptors, demonstrated in Fig. 1(c). By the target protein *X* separately regulating *S* and *R*, we can achieve more working regimes according to equation (2). Ideally, the working regime for setting a constant protein level is when *Rp* is proportional to *S*, i.e. the global feedback is the main control. Therefore the tuning performance in homogeneous population is similar to the global feedback mentioned above when applying the same control law. When some cells switch to non-contributing states or mutants appear, they often drag the circuit dynamics to a different operating point(fail-mode) where the local feedback plays more role.

We now present an example of the layered controller and obtain three different working regimes. We assume the AHL production is activated by the target protein in a linear kinetics, same as the internal feedback in equation (1) and the global feedback in equation (3). Meanwhile, the target protein regulates the receptor transcription in a Hill-type kinetics. The ODE model is obtained as the following:

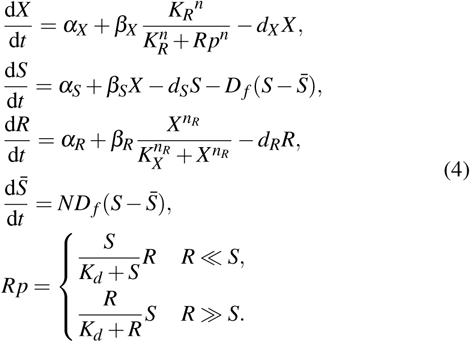

We show the tunable range of the protein level in Fig. 4(a) and working regimes of the layered controller in Fig. 4(b). Notice that when the protein level is perturbed to be lower/higher than its minimum/maximal value that the circuit can reach in the ideal linear working regime, the local feedback starts to contribute more than the global feedback in the the control actuation via *Rp*.

**Fig. 4.**
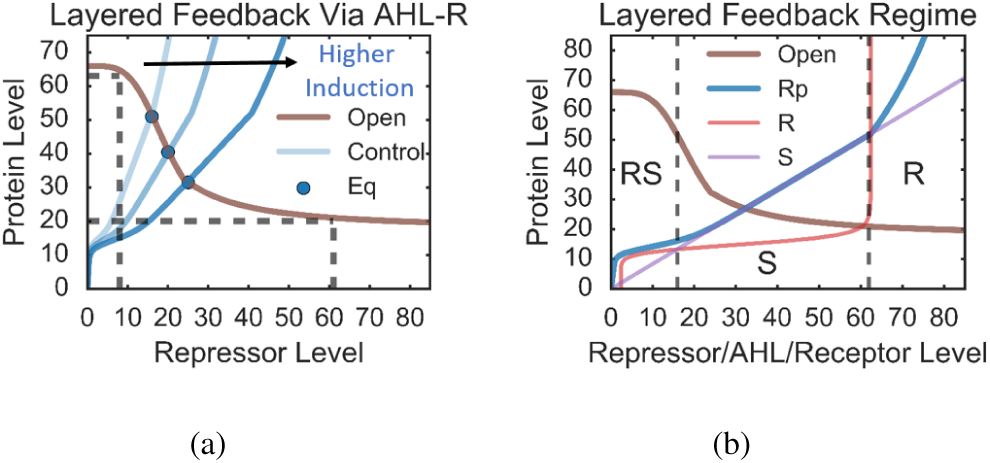
Tuning response curves of the layered feedback circuit. Panel (a) shows that the target protein level can be tuned by inducing the AHL synthesis at different strengths with a similar tuning range as the internal and the global controller. Panel (b) demonstrates three working regimes: *Rp* depending on both *R* and *S* when *X* is very low; *Rp* depending on *S* in the ideal working space; *Rp* switching to *S* saturation regime and depending on *R*.

## IV Protein Homeostasis in Heterogeneous Population

The three controllers presented above all function similarly in homogeneous populations within the ideal working regime since all cells contribute equally and follow the same response curve. However, non-contributing cells or mutants that do not work in the ideal regime might appear in populations, and the populational protein level is perturbed.

We consider two potential sub-populations that are non-contributing cells or mutants: 1) they only express a low level of protein; 2) the repression pathway on the target protein is broken. The first case often happens when the protein expression takes substantial energy and causes metabolic burden so that mutants that produce fewer proteins have more growth benefits [32]. It is also observed when cells are under starvation or shock so they switch to a low functional state with slow expression [33]. The second case can occur when the target protein offers advantage in survival thus mutants with high protein expression tend to be selected, such as antibiotic resistance [34]. Some repressors and activators are affacted by certain resources in the environment and their regulation pathways can be turned on or off as a response to environmental fluctuations [35], [36]. In this section, we show how internal, global and layered feedback controllers perform to regulate populational protein homeostasis in these two heterogeneous populations.

### A. Non-contributing Cells with Low Protein Expression

Consider the first case when non-contributing cells or mutants only express a low level of protein. To model the heterogeneous population behavior, we assume the total population consist of *N*_1_ contributing cells that are properly functional in the ideal working regime and *N*_2_ = *N* − *N*_1_ non-contributing cells in the fail-mode. We use *η* ≪ 1 to characterize how much the protein expression is slowed.

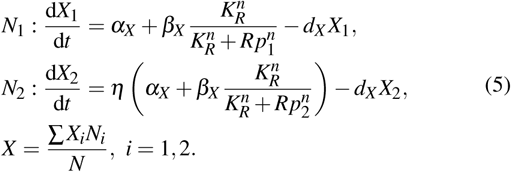

We apply internal, global and layered feedback controllers to the system in equation (5). For *i* = 1, 2 in *N*_*i*_, we obtain the following equations of repressor level *Rp*_*i*_ solved at steady state:

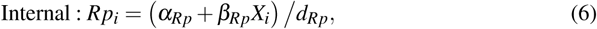

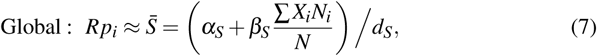

Layered:

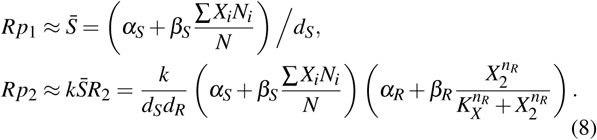

Using the internal feedback in equation (6), contributing and non-contributing cells maintain their own protein levels via the internal repressor *Rp*_1_, *Rp*_2_. When more non-contributing cells appear, the population level protein homeostasis is significantly disturbed from the normal level *X*_1_ to a lower level 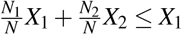 given *X*_2_ ≤ *X*_1_, as shown in Fig. 5(a).

**Fig. 5.**
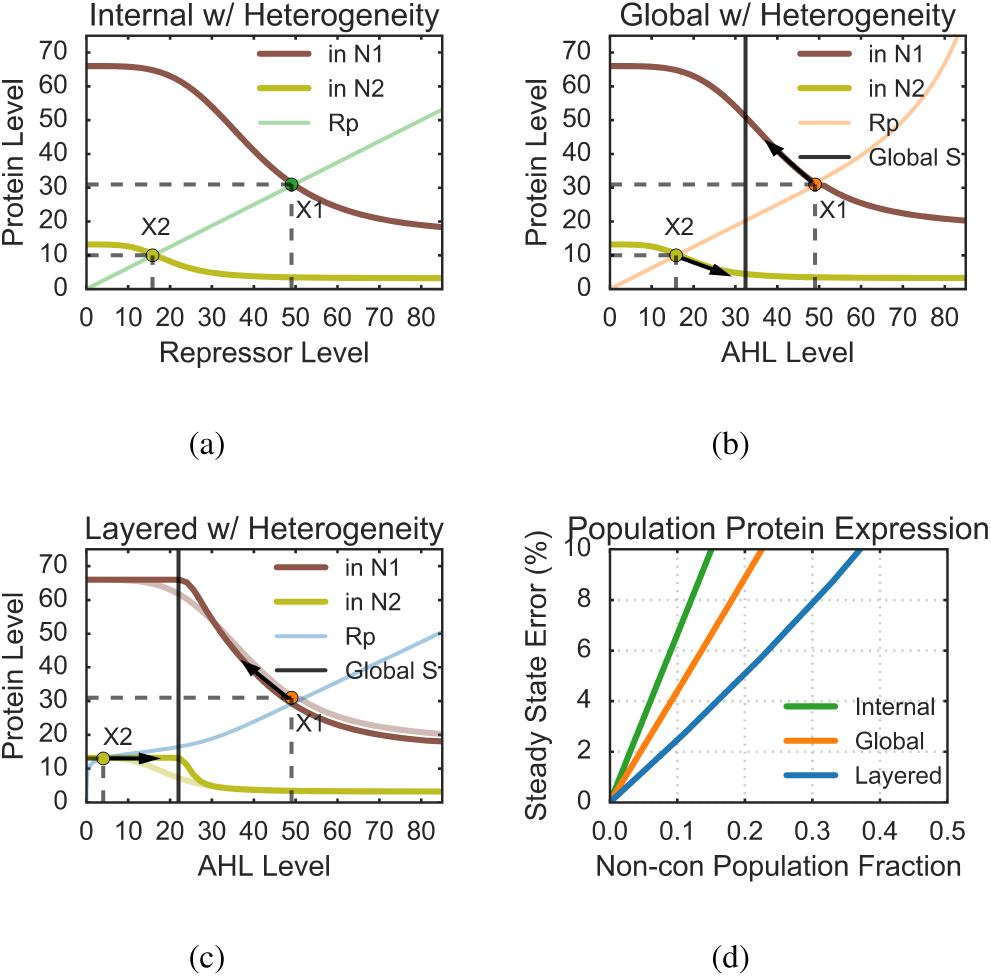
Tuning response curves and simulations of steady state error in populational protein levels across heterogeneous populations. Panel (a)-(c) are response curves of contributing and non-contributing cells using the internal, global and layered controller. In (a), the contributing cells produce an expected level of the target protein of *X*_1_, while non-contributing cells produce a low level of protein *X*_2_(the yellow dots). The population level protein expression is determined as the weighted average of *X*_1_ and *X*_2_(between the two dashed horizontal lines). In (b), the global AHL level moves to the middle(black vertical line) since it measures the populational protein level. Contributing cells are actuated to relieve the repression on *X* (black arrow pointing up from *X*_1_). The non-contributing cells also follow the global control and decrease their protein expression more, which is the opposite to the protein recovery. In (c), the local controller governs the non-contributing cells so the protein level doesn’t change much with the global feedback. The contributing cells compensate for the decrease in protein expression through the global feedback. Panel (d) compares the simulated steady state errors in population level protein expression when non-contributing cells appear. The layered controller can tolerate a higher fraction of non-contributing cells than the internal or global controller.

The global feedback shows a better control performance since both contributing and non-contributing cells share the same AHL molecules. As shown in Fig. 5(b), when non-contributing cells *N*_2_ have low protein expression, the global AHL 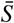 is perturbed to a lower level (the black line). Therefore, the protein expression in contributing cells *N*_1_ is steered up to a higher level to compensate and recover population level protein homeostasis (the arrow pointing out from current equilibrium). However, since the non-contributing cells are also governed by the global feedback and the control action drags the their protein expression to an even lower level, which can hinder the overall disturbance rejection across the population.

By combining the local feedback together with the global feedback, we show that the layered controller improves the robustness of protein expression in Fig. 5(c). In contributing cells *N*_1_, the global feedback governs and leads to a compensating increase in protein level. Meanwhile, the non-contributing cells *N*_2_ work in a different regime where the local controller regulates the protein level to be relative robust to variations in AHL level. Therefore, the populational protein homeostasis can be maintained by contributing cells and avoids fail-modes in non-contributing cells get worse. Notice that the tuning curve becomes a little bit sharper in the left of the plot (darker and lighter curves), but the tunability is not influenced much as the the difference appears more outside the ideal working regimes for tuning.

To obtain a more practical knowledge of how the global and the layered feedback improve the robust expression of protein across heterogeneous populations, we simulate for the steady state error of populational protein level with a range of fractions of the non-contributing sub-population. In Fig. 5(d), we compare the error versus fraction curves of three controllers, and by setting the error cap to be 10%, the maximal fraction of non-contributing cells the population can tolerate is 15% for internal, 22% for global and 36% for layered feedback. Assuming the non-contributing cells grow 20% faster than the contributing cells, applying the layered feedback can ensure the functionality for around 9 more generations than the internal feedback before the error goes beyond the limit.

### B. Non-contributing Cells with Weak Protein Repression

Similarly, we consider another scenario of heterogeneous populations where the repressor in non-contributing cells fails to inhibit the target protein’s transcription. We model such breakdown by setting a large value of *λ K*_*Rp*_, *λ* ≫ 1 in the repression Hill function and it represents a weaker repressor in binding to its DNA binding site.

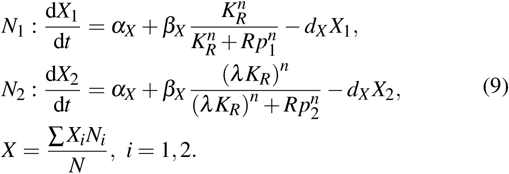

The controllers are the same as in equation (6)-(8), and we plot analytical and simulated results in Fig. 6. The internal feedback regulates contributing and non-contributing sub-populations separately and the populational protein level is the weighted average of *X*_1_, *X*_2_, shown in Fig. 6(a). The global controller via AHLs only functions within the linear regime before it saturates, and when the repression pathway stops to work, the non-contributing cells are dragged to the saturation regime. As shown in Fig. 6(b), even though the global 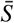 helps by providing information of population level behaviors, the control input is saturated by the constitutive receptor and cannot improve much on the disturbance response. The layered control either performs as the global feedback in contributing cells or the local feedback in non-contributing cells since the heterogeneous population work in different regimes. As shown in Fig. 6(c), the layered feedback can manage to handle the disturbance better than merely with the global controller. Fig. 6(d) demonstrates that the layered controller can maintain the steady state within 20% error for around 22 more generations than the internal feedback, assuming non-contributing cells are 20% faster in growth.

**Fig. 6.**
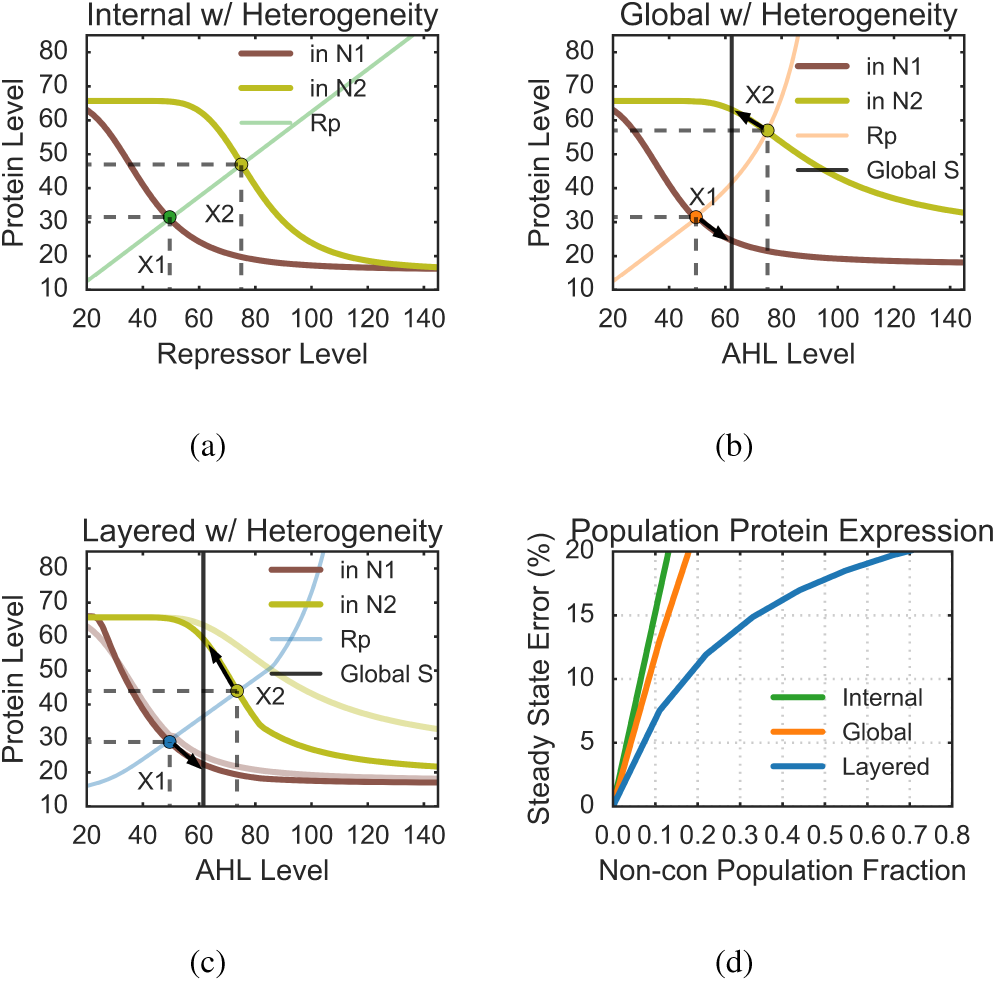
Tuning response curves and simulations of steady state error in populational protein levels across heterogeneous populations. Panel (a)-(c) are response curves of contributing and non-contributing cells using the internal, global and layered controller. In (a), the contributing cells produce an expected level of the target protein of *X*_1_, while non-contributing cells have a weaker repression on *X*, leading to a higher protein level *X*_2_(the yellow dots). In (b), the non-contributing cells has a higher protein level since the AHLs are saturated. Contributing cells and non-contributing cells are actuated by the global AHLs in opposite directions. In (c), the local controller governs the non-contributing cells and their protein level doesn’t exceed much from AHL saturation. Panel (d) compares the simulated steady state errors in population level protein expression when non-contributing cells appear.

## V. DISCUSSION

In this paper, we show that the layered feedback control structure improves the population level homeostasis in protein expression. By ODE modeling, analysis of tuning response curves and working regimes, and simulations of a simple feedback regulation circuit, we demonstrate and compare three controllers’ performances in two potential heterogeneous populations. The analyzing approach and observations from this case study tend to be applied to a more general area of population control problems. Integrating the single-cell level circuit design with cell-cell communications extends our ability to build more diverse and stable microbial consortia [37]–[39]. It also requires a new perspective in theory to understand population level behaviors, such as stability and robustness, and novel control structures across single-cell levels to multicellular levels need more investigation. In this study, we rethink the source of disturbances in cell populations and propose that layered controllers are good strategies to maintain robust functionality while having diverse population heterogeneity.

In the future, we will apply the layered control structure to more population control problems, such as controlling population size, fraction, differentiation and spatial organizations, using models of cell growth/death and states transition processes. Meanwhile, we will look into more sophisticated origins that cause population heterogeneity and connect control strategies in nature to improve synthetic designs.

## VI. ACKNOWLEDGMENTS

The authors would like to thank Fangzhou Xiao for his insightful discussion and Ronghui Zhu for the suggestion on experimental circuit design. The author X. R is supported the Defense Advanced Research Projects Agency (Agreement HR0011-17-2-0008). The content of the information does not necessarily reflect the position or the policy of the Government, and no official endorsement should be inferred.

